# Scanning probe microscopy elucidates gelation and rejuvenation of biomolecular condensates

**DOI:** 10.1101/2024.08.28.610139

**Authors:** Aida Naghilou, Oskar Armbruster, Alireza Mashaghi

## Abstract

Comprehensive understanding of dynamics and disease-associated solidification of biomolecular condensates is closely tied to analysis of their mechanical characteristics. Despite recent technical advances in rheological studies of condensates, these still vastly rely on methods restricted to small forces, rendering measurements of droplets with higher elasticities and after transition to solid challenging. Here, we develop assays for in-depth mechanical characterization of biomolecular condensates by scanning probe microscopy. We demonstrate this technique by measuring the rheological behavior of heterotypic poly-L-lysine heparin condensates, showcasing their multi-route liquid to gel transition, as well as their rejuvenation by chemical alterations to the medium. Due to the wide-spread application of scanning probe microscopy in biological fields, its capability for rapid, high throughput, high force range studies, and integration with nanoscale morphological measurements, our probe-based method is a significant breakthrough in investigating condensate behavior, leading to accelerated development of therapies.

## 1. Introduction

Biomolecular condensates have been recognized as one of the groundbreaking recent discoveries in biomedicine, unveiling a distinct organizational principle in molecular biology [1]. These are membrane-less, phase separated assemblies of proteins, nucleic acids, or carbohydrates in the form of dense liquid droplets co-existing in the liquid environment of cells, extracellularly, or in a synthetic liquid surrounding [2, 3]. Cellular structures such as nucleoli, paraspeckles, Cajal bodies, P-bodies, germ granules, and stress granules are examples of biomolecular condensates [4-7]. They mediate a variety of cellular functions ranging from stress response to transcription by rapid assembly and disassembly or cellular signaling events [8, 9]. The high local concentration within the dense phase of condensates has the potential to enhance aberrant protein interactions, leading to aggregation [10, 11]. This improper assembly can lead to solidification, causing conformational diseases such as amyotrophic lateral sclerosis, and neurodegenerative disorders where amyloid fibril formation is a hallmark of pathology [12-15].

While mechanical processes are key to condensate related pathology and therapeutic developments, assays suitable for directly probing liquid-solid transition are lacking. Rheological methods are commonly used to demonstrate the liquid behavior of biomolecular condensates, and to investigate the nature of molecular interactions and droplet dynamics [10, 16]. The active microrheological characterization of phase separated condensates is mostly performed by means of optical tweezers (OT) [8, 16-20]. While this technique enables measurements with very high force resolution and precision, the force range available with OT is limited, which constrains measuring the mechanics of droplets with higher elasticities and after fibril formation [21-23]. With the increasing discoveries of nondynamic, solid-like condensates [24], novel strategies compatible with their mechanics are crucial. Furthermore, due to the high laser powers, OT is often not suitable for studying condensates composed of thermally sensitive biomolecules [25]. A notable limitation of OT in rheological studies of biomolecular condensates is the necessity of introducing beads to the system for trapping and manipulation. Even at low concentrations, these may interact with condensate components, triggering biochemical responses that are not well characterized [26]. This may potentially alter the mechanical behavior of condensates, even without significantly impacting their assembly and morphology. In addition to OT, micropipette aspiration has also been used for assessing the mechanics of condensates [10, 27-29]. This method is, however, limited to condensates with characteristics of Newtonian fluids, assumes a homogeneous system, cannot be used on amyloid fibrils, and is challenging to use at higher frequencies due to inertia effect during micropipette aspiration [22, 27, 29]. Fluorescence recovery after photobleaching (FRAP) is a widely employed technique, providing information on condensate fluidity, viscosity, and diffusion [8, 30-36]. However, it does not provide information on the interfacial mechanics and is challenging to interpret for multi-component droplets with diverse velocities and specifically for condensates with static behaviors [22, 25]. In addition, it has been shown that fluorescent labeling of proteins may alter the properties of condensates [37].

Given the aforementioned limitations, developing new characterization methods for quantifying condensate mechanics is invaluable. In particular, there is a need for a technique that enables single droplet mechanical measurements with the same device from liquid to solid, as using various methods may lead to results that at times differ by several orders of magnitude [38]. One strategy that has gained interest in mechanobiology is the use of scanning probe microscopy (SPM) and its variant atomic force microscopy (AFM). The main advantages of AFM over OT are a wider force range, ubiquity in various biological fields, as well as combining force measurements with nanoscale morphological investigations. Due to the much higher forces exerted by AFM probes, it has been successfully employed to measure the mechanical properties of amyloid fibrils [23, 39-44]. Therefore, probe-based techniques such as SPM and AFM are great candidates for assessing the full range of condensate mechanics from liquid to fibril using the same device, *in situ*. Recently, studies have explored quantifying the ultra-low interfacial tension of coacervates by colloidal-probe AFM [22, 25, 45]. However, these studies were limited to measurements of interfacial energy of capillary bridges. A comprehensive characterization of elastic and viscous shear moduli of single condensate droplets, akin to what has been achieved with OT has not yet been realized with AFM.

In addition, biomolecular condensates are scaffolded by fibrillar networks and dynamic environments within a cell, which changes their physical properties such as coalescence and morphologies [46-48]. This, together with the growing interest in wetting of membranes by liquid droplets, calls for the development of approaches that allow measuring the mechanics of condensates under indentation forces, enabling a better understanding of interface interactions.

In this work, we address these gaps in the field of condensate biology by developing an innovative approach for studying condensate mechanics by means of SPM. To demonstrate this methodology, we employ heterotypic condensates of Poly-L-lysine (pK) and heparin (H), as a model system, subjecting them to indentation and deformation at various frequencies, and develop a data analysis approach to interpret the results. We study the rheological properties of condensates and demonstrate their multi-route liquid to gel transition, as well as their rejuvenation. Given the common use of SPM in biomedical research, its ability to provide rapid, high-force range measurements at high throughput, and its integration of force measurements with nanoscale morphological analysis, our SPM-based assay offers a unique advantage in characterizing liquid-solid transitions in condensates.

## 2. Results

Our workflow involved selecting a condensate from phase contrast images (Figure 1A, supplementary video 1), landing the probe on the droplet, indenting it with a preload, and subsequently modulating the indentation at various frequencies. The recorded data consists of the relative SPM head height (vs. an arbitrary zero point) and cantilever deflection force over time, *h*_r_(*t*) and *F*(*t*), respectively. A representative curve of *F*(*t*) is depicted in Figure 1B, the corresponding *h*_r_(*t*) can be found in supplementary Figure S 1A. Figure 1B shows the approach segment, where the head is lowered at a constant rate until a predefined deflection is reached (in gray), a resting period to ensure equilibrium (in red), constant amplitude head height oscillations at various frequencies (each frequency separately colored), and finally the retraction (in yellow). The pause and retraction segments are not further used in the evaluation.

**Figure 1:**
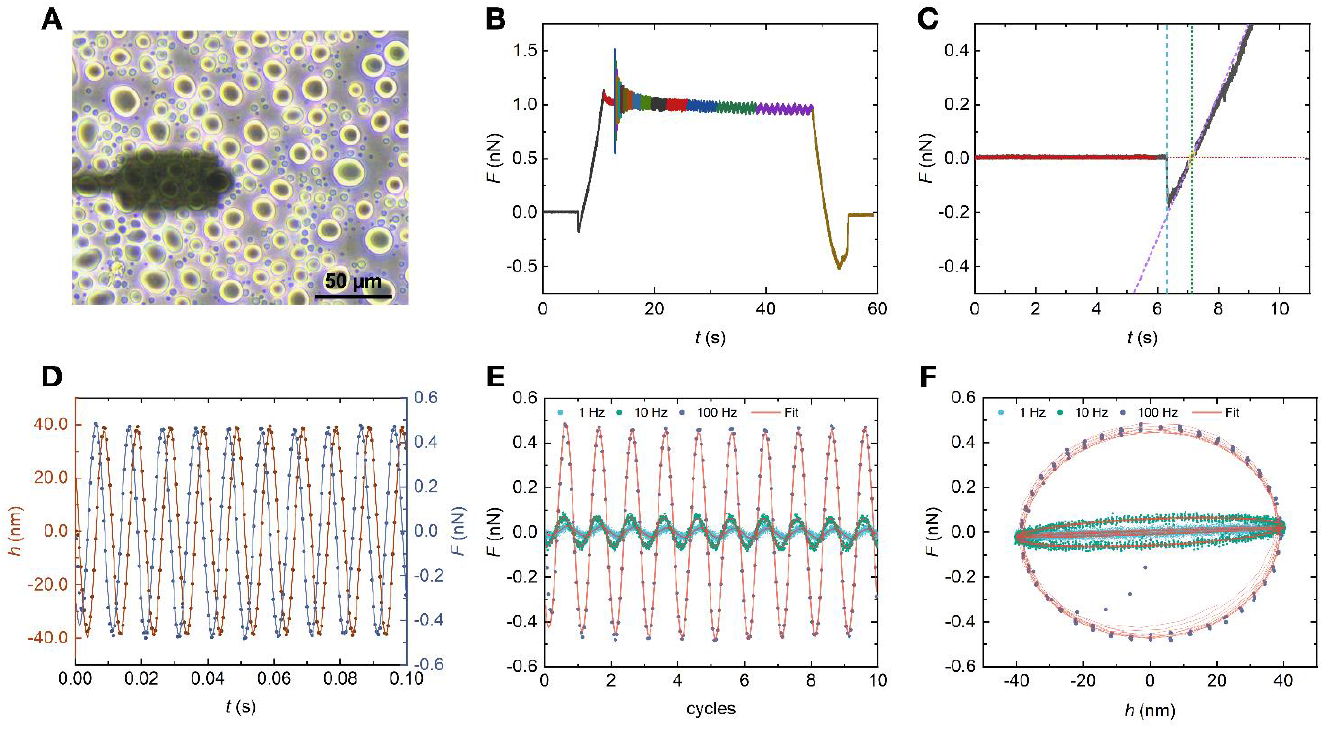
Head height and cantilever deflection force traces of condensates. **A)** Representative phase contrast micrograph depicting the top view of condensates and the indenter resting on a droplet. Scale bar is 50 µm. **B)** A representative curve of cantilever deflection force over time, *F(t)* showing the approach in gray, pause in red, oscillations at various frequencies separately colored, and retraction in yellow. **C)** Magnification of the approach segment of the *F(t)* curve with the baseline in red, time point of contact in cyan, approximate intersection data points in yellow, intersection fits in magenta and green. **D)** Representative experimental head height *h(t)* and cantilever deflection force *F(t)* over time and the corresponding fits with Eq. 1 and 2 for 100 Hz. **E)** Exemplary *F(t)* curves within 10 cycles of oscillatory droplet deformations for 1, 10, and 100 Hz, showing increased force at higher frequency. **F)** Exemplary force-indentation Lissajous curve at 1, 10, and 100 Hz, demonstrating higher force and phase shift with frequency.

### 2.1. Indentation determination

Figure 1C shows the deflection signal of a typical approach for the liquid droplets. As the cantilever approaches the surface, at a certain distance an abrupt contact between the indenter and the droplet occurs, which is detectible as a sharp drop in force, i.e., the cantilever is pulled towards the condensate. This is followed by a reduction of the (negative) force until the particle returns to the unperturbed geometry near zero force as the cantilever continues to approach the substrate. After the zero force is re-reached, positive forces correspond to the indentation of the sample from the unperturbed shape. To accurately evaluate the true indentation of undistorted droplets from the approach part of the curves, firstly, a moving average filter over 100 points was applied to the numeric values to reduce noise and allow a better detection of the abrupt contact point. Afterwards, the numerical derivative of the signal was computed which allows the sudden change of force to appear as a peak in the derivative data (supplementary Figure S 1B). This leads to better determination of the moment of contact depicted as cyan vertical line in Figure 1C. With a short keep-out period, the data prior to the contact point (red in Figure 1C) are linearly fitted to determine the baseline and are extrapolated to the approximate intersection time. A region of ± 200 ms shown as yellow data points around the approximate intersection is linearly fitted (magenta line) and the exact intersection of the two fitted curves is considered as the time of zero indentation *t*_0_ (green vertical line) and the head height of zero indentation can be read from *h*_*r*_(*t*_0_) = *h*_0_. With *h*_0_, the relative head height data can be corrected with *h*(*t*) = *h*_r_(*t*) – *h*_0_ to accurately reflect the true time-dependent indentation of the condensates.

#### 2.2. Droplet mechanics

After the indentation and the pause period, the head height was harmonically driven for 10 cycles at each frequency, leading to an indentation modulation of the condensates. Each oscillation frequency was evaluated separately and both *h*(*t*) and *F*(*t*) were fitted with

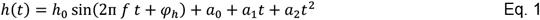

and

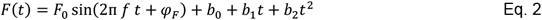

consisting of a sinusoidal function of frequency, *f* and phase φ, on a polynomial baseline to account for slight signal drift during the measurement, which may originate form e.g., the temperature drift of the cantilever or the creep of the droplets. Hence, the indentation depth δ varies with time and is calculated separately for each frequency as the time average of *h*(*t*) (Supplementary Figure S 2). Exemplary *h*(*t*) and the resulting *F*(*t*) are shown in Figure 1D for 100 Hz, where the phase between the force and height data is visible. Supplementary Figure S 3 allows a comparison of the fits for various frequencies and hints at the strong frequency-dependent response of the liquid droplets observable by the larger forces at higher frequencies at constant head height amplitudes, which is also visible in Figure 1E and F.

To determine the complex frequency dependent rheological behavior from the measured data, it is necessary to consider the contribution of the cantilever and the surface energy (Figure 2A). From *F*_0_ and *h*_0_, and their corresponding phases, φ_F_ and φ_h_, the complex experimental spring constant *X*_e_ is calculated as

**Figure 2:**
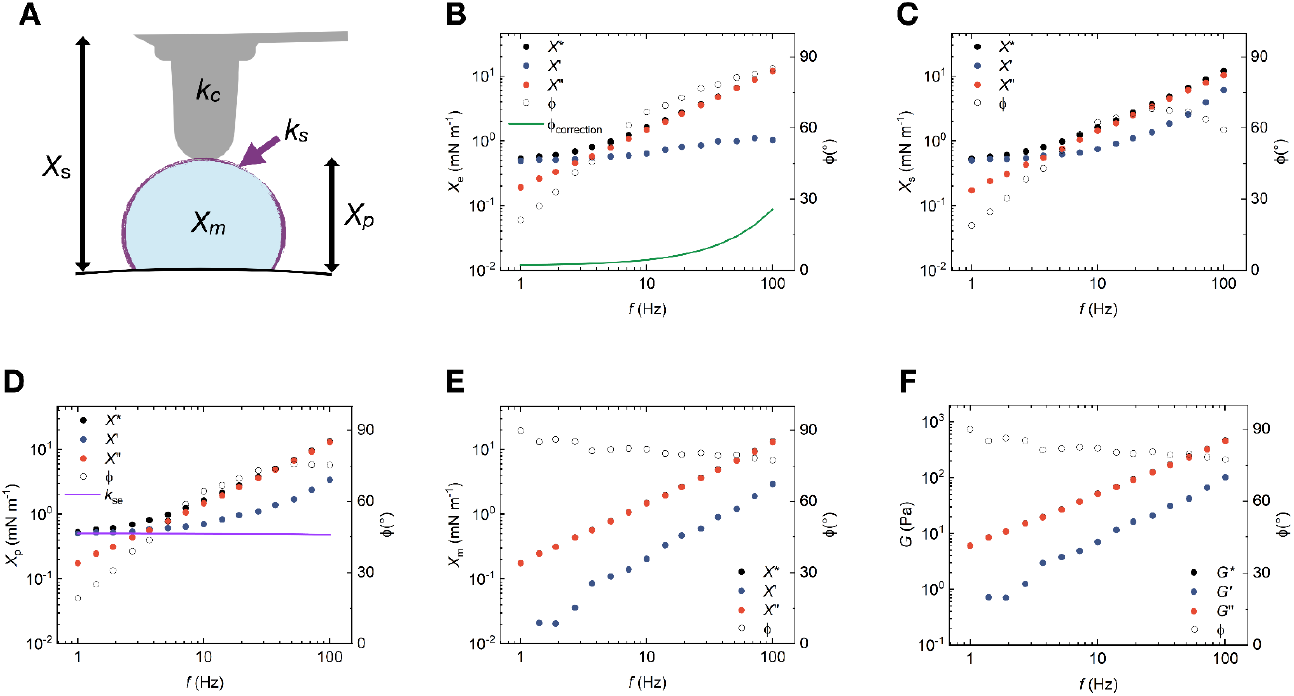
Evaluation of condensate complex shear moduli. **A)** Schematic of the contribution of multiple spring constant to measurement, *X*_*s*_ : complex system spring constant, *X*_*p*_: complex particle spring constant, *X*_*m*_: complex material spring constant, *K*_*c*_: cantilever spring constant, *k*_*s*_: surface energy as a spring constant. **B)** Complex experimental spring constant *X*_*e*_ calculated from Eq. 1-3, and the corresponding real *X’*_*e*_ and imaginary *X”*_*e*_ contributions, as well as the corresponding phase φ, and phase lag φ_correction_ calculated from Eq. 7 and 8 (see Methods). **C)** Complex system spring constant *X*_*s*_ calculated from Eq. 4, and the corresponding real *X’*_*s*_ and imaginary *X”*_*s*_ contributions, as well as the corresponding phase φ. **D)** Complex particle spring constant *X*_*p*_ calculated from Eq. 5, and the corresponding real *X’*_*p*_ and imaginary *X”*_*p*_ contributions, as well as the corresponding phase φ and the surface energy spring *k*_se_ calculated from Eq. 9 and 10 (see Methods). **E)** Complex material spring constant *X*_*m*_ calculated from Eq. 6, and the corresponding real *X’*_*m*_ and imaginary *X”*_*m*_ contributions, as well as the corresponding phase φ. **F)** Complex shear modulus G calculated from Eq. 11-14 (see Methods), and the corresponding real (elastic modulus, *G’*) and imaginary (viscous modulus, *G”*) contributions, as well as the corresponding phase φ.

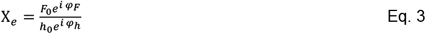

For the data shown in Figure 1B and utilizing Eq. 3, *X*_e_ and the corresponding phase as well as its real and the imaginary parts (*X’*_e_ and *X’’*_e_, respectively) are deduced and depicted in Figure 2B.

*X*_e_ still contains a phase lag caused by hydrodynamic drag of the surrounding fluid on the cantilever as depicted by the green line in Figure 2B. Compensating the phase lag (see Methods) leads to the system spring constant *X*_s_ (Figure 2C) with

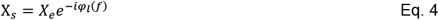

*X*_s_ is itself a series connection of the cantilever spring *k*_c_ and the particle spring *X*_p_. Thus, *X*_p_ (Figure 2D) can be calculated as

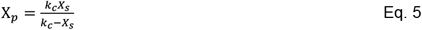

*X*_p_ is composed of a connection of springs, in this case a parallel connection of a spring resulting from the surface energy change due to deformation of the particle *k*_se_ (see Methods) [16, 49] and the spring resulting from the bulk deformation of the viscoelastic material *X*_m_. *k*_se_ can be determined from the low-frequency value of *X*_p_’, which is shown as the magenta line in Figure 2D. Hence, *X*_m_ (Figure 2E) is given by

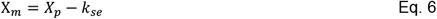

This results in *X*_m_’= 0 at the frequency utilized for calculating *k*_se_. Hence, all values of *X*_m_’ at and below this frequency are excluded from further evaluations [16, 49].

Using the Hertzian contact model (see Methods) the complex shear modulus of the material, *G* and the corresponding real (elastic modulus, *G*’) and imaginary (viscous modulus, *G*”) contributions can be determined (Figure 2F). A linear fit of *G*” over *f* leads to the determination of the viscosity, η of droplets (Supplementary Figure S 4).

### 2.3. pK-H condensate properties

Biomolecular condensates interact with their environment, which causes changes in their properties on a molecular level leading to variations in rheological behavior [27, 50-52]. It is well-known that the rheological characteristics of condensates is salt-dependent and condensate viscosity typically decreases when droplets are formed at higher salt concentrations [16, 32, 49, 50, 53]. Using our SPM-based technique and evaluation, the material properties of pK-H condensates were evaluated. Condensates formed at KCl concentrations between 0.85 and 1.1 M were probed to determine *G*’, *G*”, and η. In addition, η was determined from FRAP experiments in parallel to compare the results with SPM.

The rheological characteristics of the condensates is calculated employing Eq. 1-6 and is provided in Figure 3A-D. Droplets at all studied concentrations show a dominant viscous behavior at the experimentally accessible frequency range. η was found to be (6.9 ± 0.2) Pa s at the lowest concentration of 0.85 M and decreased to (5.2 ± 0.2), (4.9 ± 0.2), and (2.29 ± 0.04) Pa s for 0.9, 1, and 1.1 M, respectively. Similar trends were observed in FRAP experiments, where a faster fluorescence restoration at higher salt concentrations is evident from the smaller values of the half-time of recovery, τ_1/2_ and larger diffusion coefficients, *D* (Figure 3E). Representative confocal images of droplets recovery within 2 min are exemplified in Figure 3F. Using Eq. 15-18, η was calculated form FRAP and compared with SMP results as shown in Figure 3G. η values from FRAP and SPM are in good agreement, validating our newly developed approach.

**Figure 3:**
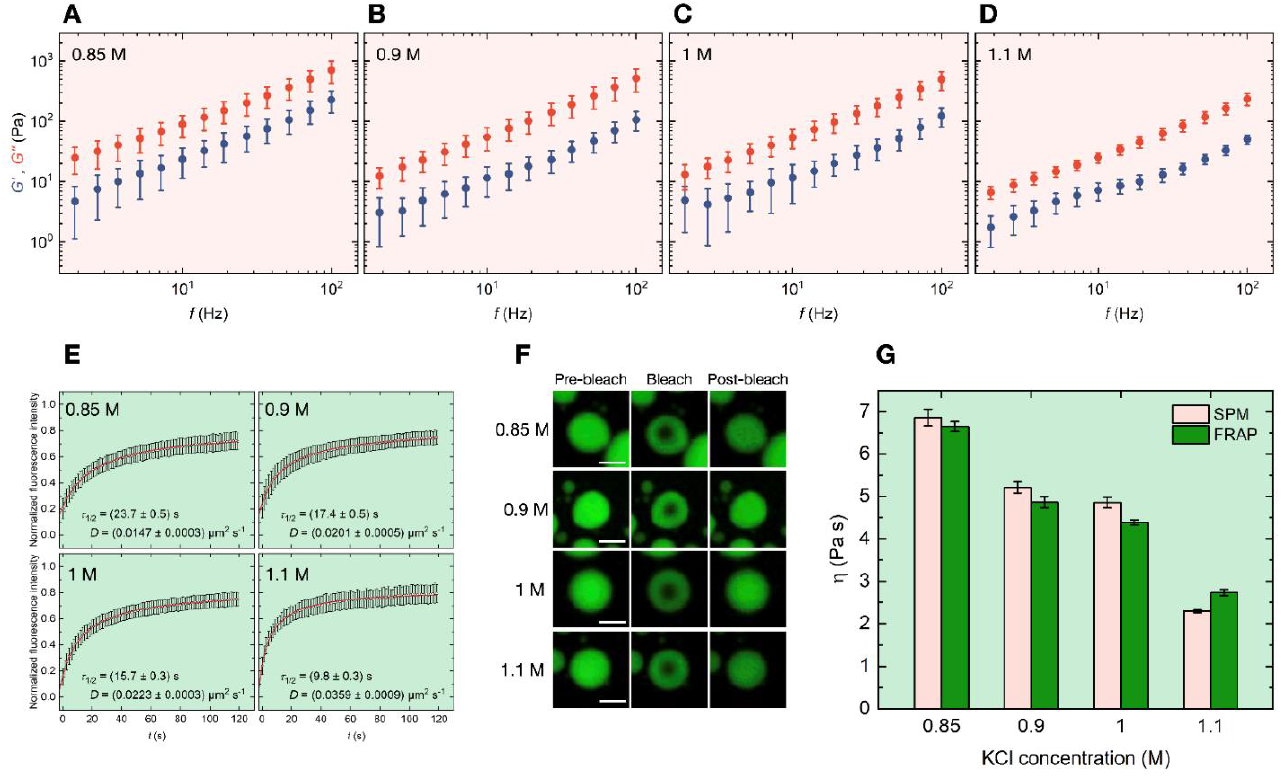
Salt dependent rheological behavior of pK-H condensates. Elastic (*G*’, blue) and viscous (*G*”, red) shear moduli of pK-H condensates formed at **A)** 0.85 M KCl (mean ± SD, *N* = 21 from three independent sample preparations). **B)** 0.9 M KCl (mean ± SD, *N* = 29 from three independent sample preparations). **C)** 1 M KCl (mean ± SD, *N* = 19 from three independent sample preparations). **D)** 1.1 M KCl (mean ± SD, *N* = 27 from three independent sample preparations). **E)** Time-dependent normalized fluorescence intensity depicting the recovery of the bleached area (circular data points) as well as the fit with Eq. 16, resulting in half-time of recovery (τ_1/2_) and diffusion constant (*D*) with Eq. 17 for each salt concentration (mean ± SD, *N* = 23 from three independent sample preparations). **F)** Representative confocal micrographs of droplet recovery after photobleaching, scale bar in all micrographs is 10 µm. **G)** Comparison of viscosity η calculated form the linear fit of *G*” over *f* in A-D measured with SPM (pink) as well as from the FRAP measurements (green) calculated with Eq. 18.

Currently, the mechanical properties of condensates is evaluated by means of methods that rely on their liquid nature [54], rendering the measurements of transition to solid and fibril form unachievable. In order to demonstrate the applicability of SPM for measuring condensate mechanics from liquid to gel, pK-H droplets were prepared at 1 M KCl and let to settle until most coalescence events ceased. Afterwards, the supernatant was diluted to 0.65, and 0.5 M KCl and the same droplets were measured at all concentrations. At 1 M, the liquid nature of the condensates is visible from their morphology and rheological characteristics as seen in Figure 4A and B, respectively. Adjusting the supernatant to 0.65 M induced a transition in the rheological properties from mainly viscous towards elastic as seen in Supplementary Figure S 5A. Further dilution to 0.5 M induced a clear alteration in the morphological and rheological properties of the condensates. Visually, they transform form liquid-like and circular, to heterogenous, rigid looking droplets with irregular borders (Figure 4A). This is in-line with the shift of the mechanical properties form liquid to gel as evident by the dominant *G*’ in Figure 4C and the frequency independence *G*’ and *G*” [55-57]. Interestingly, with gelation of condensates the approach segment of the force curves also changed from the abrupt contact seen for liquid droplets in Figure 1B, to a linear rise of *F*(*t*) as seen in Supplementary Figure S 6 and was evaluated differently from the liquid condensates (Supplementary Information).

**Figure 4:**
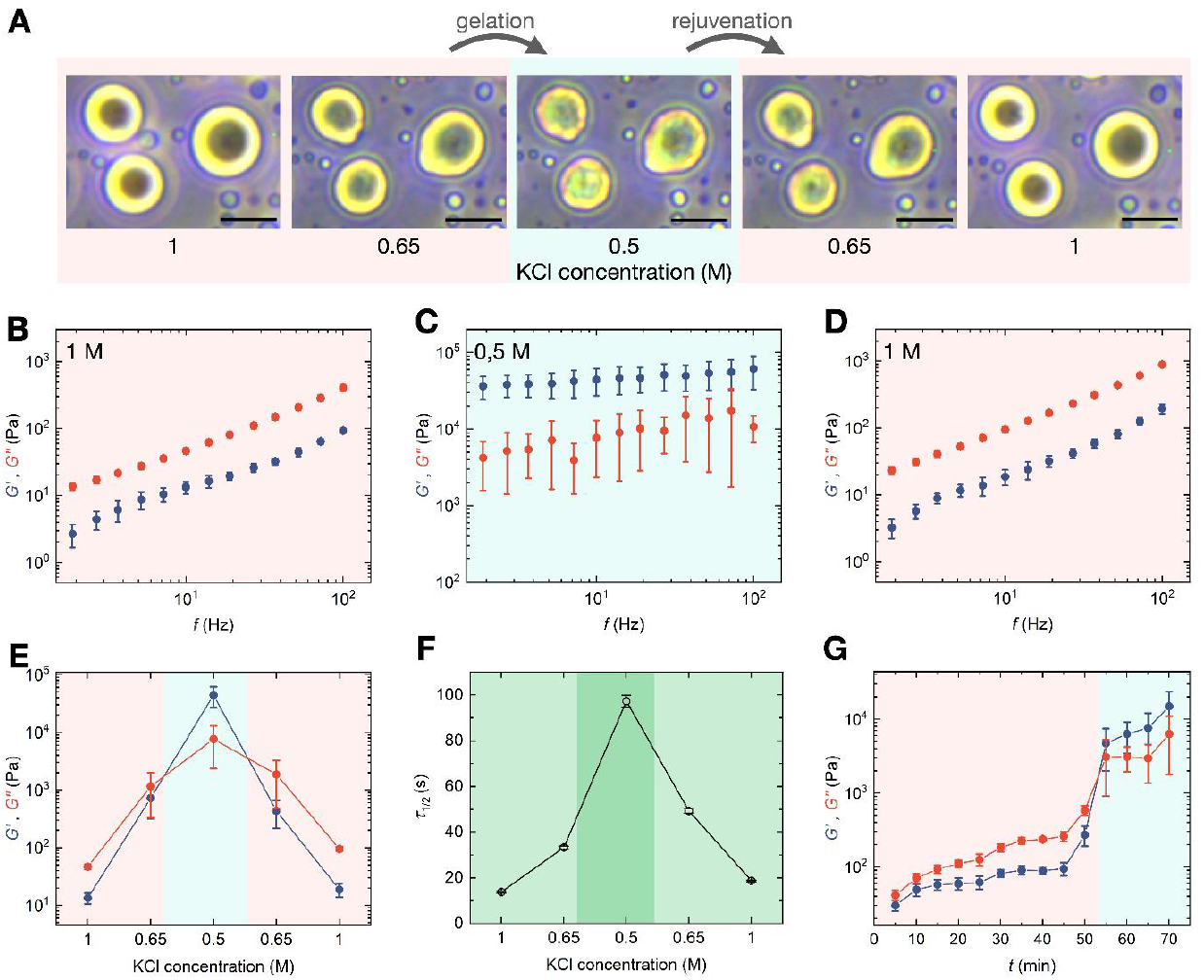
Gelation and rejuvenation of pK-H droplets. **A)** Representative phase contrast micrographs of droplets formed at 1 M, and after diluting the supernatant to 0.65 and 0.5 M KCl to induce gelation as well as exchanging back to 0.65 and 1 M to trigger rejuvenation, scale bar in all micrographs is 10 µm. **B)** Elastic (*G*’, blue) and viscous (*G*”, red) moduli of pK-H condensates formed at 1 M KCl and **B)** Measured at 1 M KCl (mean ± SD, *N* = 10). **C)** Measured after dilution of supernatant to 0.5 M KCl (mean ± SD, *N* = 6). **D)** Measured after exchange of supernatant back to 1 M KCl (mean ± SD, *N* = 10). **E)** *G*’ and *G*” of droplets at 10 Hz displaying the transformation of condensates from liquid to gel-like by dilution of supernatant and a subsequent rejuvenation after the supernatant was reverted to 1 M KCl. **F)** Half-time of recovery τ_1/2_ highlighting the slower fluorescence restoration for gel-like condensates and faster after rejuvenation (mean ± SD, *N* = 10). **G)** *G*’ and *G*” of droplets at 10 Hz depicting the temporally resolved conversion of condensates from dominantly viscous to elastic (mean ± SD, *N* = 10).

As the ideal method for characterizing condensates would be capable of assessing both solidification and rejuvenation, the alteration of the condensates based on their response to salt concentration of the medium prompted us to investigate whether this behavior is reversible. Hence, we subsequently exchanged the KCl concentration from 0.5 M back to 0.65 M, which returned the condensates from gel-like to a transition state between elastic and viscous characteristics (Supplementary Figure S 5B). Returning to 1 M KCl medium also recovered the liquid state of droplets and their vastly viscous behavior (Figure 4D). This rejuvenation is also evident by the morphological changes of droplets as depicted in Figure 4A and Supplementary video 2.

Figure 4E depicts *G*’ and *G*” at 10 Hz during the transition from liquid to gel and the following rejuvenation. Both the lines and the color-coded background are a guide for the eye. A hysteresis of the mechanical properties can be detected by the higher values of *G* after rejuvenation, which might be related to structural rearrangements within the material. FRAP experiments (Figure 4F) performed on the condensates also demonstrated significantly higher τ_1/2_ values at 0.65 and 0.5 M KCl, with a return to faster recoveries after rejuvenation with a similar hysteresis as observed by SPM mechanical evaluations. The color-coded background resembles the SPM measurements. The corresponding recovery traces and *D* at various concentrations can be found in Supplementary Figure S 7.

An important aspect of developing methods to measure condensate mechanics from liquid to solid state is the ability of the device to perform *in situ* assessment of droplets. To demonstrate liquid to gel transition *in situ* and temporally resolved, we increased the humidity in the measurement chamber, by heating up deionized water to 55°C and letting the vapor incorporate into the dilute phase, decreasing the KCl concentration. *G*’ and *G*” at 10 Hz are depicted in Figure 4G, where a slow adsorption of humidity caused a gradual increase in both moduli until enough dilution rendered the condensates to obtain gel-like behavior with prevailing elastic properties (Figure 4G).

To show the versatility of our method, we next applied it to study gelation of droplets induced by chemical crosslinking. We employed a well-stablished chemical crosslinking agent, formaldehyde (FA), to pK-H droplets to determine whether it can induce gelation. Despite its wide use in biology as a fixative, FA for fixating biomolecular condensate has not been investigated yet. Therefore, we added FA to the condensates with an end concentration of 4%, which is routinely used to fix biological specimens. As a control (CTRL) experiment, 4% water was added to the condensates. In both samples, the supernatant was exchanged with the supernatant of another drop to verify that gelation is independent of dilute phase perturbation. Indeed, the CTRL experiment demonstrates liquid-like rheological behavior (Figure 5A), while the crosslinked droplets have gel-like characteristics (Figure 5B). This is also validated by striking differences in the diffusion within the droplets in FRAP experiments, where there is hardly any recovery for FA-treaded condensates (Figure 5C), while the CTRL droplets show much faster fluorescence restoration evident by the smaller values of τ_1/2_ and nearly an order of magnitude higher *D* (Figure 5D).

**Figure 5:**
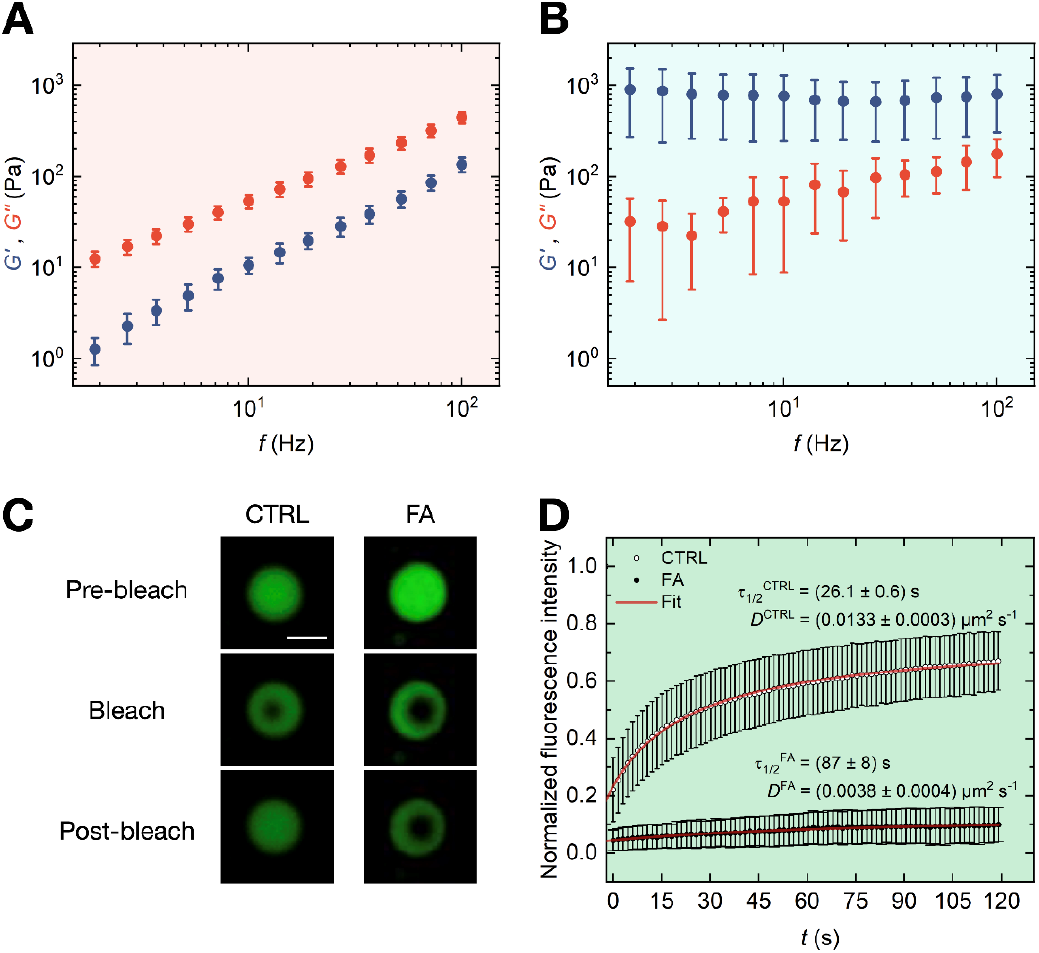
Crosslinking of pK-H droplets. Elastic (*G*’, blue) and viscous (*G*”, red) shear moduli of pK-H condensates formed at 1 M KCl **A)** After addition of water to a concentration of 4% for control (CTRL) experiment (mean ± SD, *N* = 12). **B)** After addition of formaldehyde (FA) to a concentration of 4% (mean ± SD, *N* = 8). **C)** Representative confocal micrographs of droplets depicting the fluorescence recovery after photobleaching for CTRL experiments and a lack of recovery after crosslinking for FA-treated condensates. **D)** Time-dependent normalized fluorescence intensity (round data points) as well as the fit with Eq. 16, resulting in half-time of recovery τ_1/2_ and diffusion constant *D* with Eq. 17 (mean ± SD, *N* = 17 for CTRL, *N* = 10 for FA).

## 3. Discussion

Overall, with the rising interest in quantifying the material properties of liquid droplets, the advancement of novel characterization techniques becomes more essential [54]. In this study, we established an approach that allows measurement of the full range of rheological properties of single microdroplets under indentation forces, a geometry that is relevant in the cellular context, where they are confined and in contact with other membranes and surfaces [50, 58]. This is increasingly growing in significance due to interest in the membrane wetting by liquid droplets [27, 59-61]. Our approach, which allows measuring the mechanical forces exerted by two interfaces on each other, has the potential to greatly advance the understanding of condensate-membrane interactions by enabling novel *in vitro* models, monitoring of the biophysicochemical properties. Another key motivation for developing this methodology was to allow measuring the rheological behavior of condensates with SPM, which has been used for mechanical testing of amyloid fibrils [39-43]. Currently, the mechanical properties of liquid droplets are typically measured by bead-based techniques such as particle tracking, oscillating beads within the condensate, or by stretching them between two beads [8, 16, 20, 49]. However, these approaches are not suitable for solid samples, where indentation experiments are the primary technique of use. Therefore, we aimed to develop a method that applies the same indentation principle on liquid droplets.

While investigating the mechanical behavior of liquid droplets, in view of their length scale, it is crucial to describe the interfacial mechanics as well as bulk viscoelasticity, ideally simultaneously measuring both [22, 54]. We showed that for determination of the frequency-dependent complex shear modulus and the interfacial surface tension of biomolecular condensates with SPM, many aspects of the system need to be considered. These include the compensation of hydrodynamic drag on the cantilever, calculating the complex system spring constant, quantifying the contribution of cantilever spring as well as the interfacial surface energy due to droplet deformation, and finally utilizing the contact geometry between the probe and condensate for the determination of the complex shear modulus based on Hertzian contact mechanics. These distinct approaches lead to our markedly optimized method, superior to previous attempts with colloidal force AFM, where the interfacial energy of capillary bridges formed from coacervates were measured [22, 62, 63]. In these studies, specific segregation proteins had to be used with the primary requirement of preferential wetting on the probe in order to form capillary condensation bridges between the probe and substrate, greatly restricting the prevalence use of this method for other phase separated solutions. In addition, the pull-off forces measured on the capillary bridges can only be attributed to interfacial mechanics. By incorporating the Hertzian contact model and representing surface energy as a spring, we could successfully attain an in-depth characterization of the rheological behavior of condensates, regardless of their nature, with details previously achieved by OT.

Our results demonstrate the applicability of SPM for precise measurement of biomolecular condensates and resolving even minor changes in their material properties by adjusting the salt concentration. The viscosities measured by SPM were found to be in good agreement with the established method of FRAP. It is important to note that similar to other biomaterials, the rheological properties of biomolecular condensates can vary depending on the measurement method.

An example is the work by Mangiarotti *et al*. [50], which reported condensate viscosities on the order of several hundred to several thousand Pa s measured by plate rheology, which are orders of magnitude higher than commonly reported values of single Pa s measured by OT [8, 16, 20, 49]. Although the viscosities obtained in our study with SPM are within similar range as OT, they are slightly higher than those obtained for pK-H previously measured by Ghosh *et al*. [8]. The quantitative agreement between our SPM and FRAP data suggests that the discrepancy between SPM and OT can be attributed to differences in preparation protocols and the experimental setup, particularly since condensates are pre-deformed before modulation. Furthermore, the multiaxial nature of forces exerted by SPM may lead to differences between mechanical properties measured by SPM and OT as previously seen in cell studies [38]. These observations highlight the potential of SPM-based methods and underscore the need for future development to investigate their impact on liquid droplets, given that condensates experience multiple forces and preloads inside the cell.

One important aspect of the presented methodology is its aptitude for investigating both reversible and irreversible gelation processes. This capability is especially valuable in medical and pharmaceutical research, as it can be used to detect pathological gelation and reveal the effects of drugs designed to reverse gelation. Our assay allowed us to vary the material properties of phase separated droplets, by exchanging the medium, firstly to lower salt concentrations inducing a transition to gel and a following increase of salt concentration rendering the droplets to rejuvenate to liquid form. We also demonstrated that irreversible gelation can be characterized by FA treatment of condensates. According to our knowledge, this is the first time that the mechanical properties of biomolecular condensates have been studied from liquid to gel and after rejuvenation. Preventing and reversing aberrant phase transitions by modulating biomolecular condensates is the ultimate goal of research targeting condensate-related pathologies [64-67]. In-depth knowledge of condensate behavior in normal and aberrant states, and after rejuvenation paves the way towards providing new understandings of diseases and enabling novel therapeutic opportunities. Hence, our study grants a versatile tool which has the potential to significantly advance both materials research and drug discovery by delivering insights into the mechanics of condensates during solidification and rejuvenation.

## 4. Methods

### 4.1. Droplet preparation

All experiments were performed on virgin polystyrene culture dishes and all chemicals were purchased from Sigma-Aldrich unless otherwise specified. The dishes were passivated with 1 w/v% BSA (A2153) solution incubated for 10 min and washed 5-6 times using deionized water. The working buffer for the induction of coacervation consisted of 1 M KCl (p9333), 10 mM imidazole (56750), and 1 g/l NaN_3_ (8.22335) [8]. The crowding agent Ficoll^®^ PM 70 (F2878) was added to the buffer with a concentration of 50 g/l to mimic the environment of a cell [68]. Afterwards, firstly heparin (H3393) and then pK (P2658) were added to reach a concentration of 40 µM each. For the salt concentration dependency experiments, all other concentrations were kept as stated above and KCl was varied between 0.5 and 1.1 M.

### 4.2. SPM measurements

A JPK CellHesion 200 (Bruker, Germany) equipped with a phase contract microscope delivering real-time *in-situ* images of measurements by means of a 20X/0.4 objective, was employed in this study. A 40 µl bulk drop containing the condensates was prepared and the droplets were let to settle on the dish. All experiments were performed within 2 h of droplet formation [49]. Afterwards, a 5 µl drop was carefully taken from the bulk liquid and placed on the cantilever to avoid the generation of air bubbles when the head is placed in and at the same time to not disturb the concentration in the dilute phase. Thereafter, the SPM head was placed inside the bulk drop, generating a capillary and setting the cantilever under the liquid. The cantilever holder is enveloped by a silicone ring, closing the sample chamber after the head is placed. To ensure constant humidity inside the chamber and inhibit condensate evaporation, a saturated potassium sulfate aqueous solution was added to the chamber, keeping the humidity at 95-97% in the temperature range of 25 to 55°C [69]. As it has been shown that micrometer-scale indenters lead to a better signal to noise ratio in comparison to their nano-scale counterparts [70], for all experiments SAA-SPH-5UM cantilevers (Bruker, Germany) made of Si_3_N_4_ with a hemispheric tip (23 µm height, 5.13 µm radius) were used. The calibration of the amplitude and the exact cantilever spring constant was determined with the thermal noise method [71]. The cantilever was passivated with 1% Pluronic (P2443) for 30 min to prevent the adhesion of droplets on the tip after measurements. This coating was specifically chosen as it has been shown to be suitable material to passivate hydrophobic surfaces and avoid unspecific interactions, as well as changes to the properties of condensates [4, 72]. Condensates were indented with forces between 0.2 and 0.5 nN and a velocity of 1 µm s^-1^, and amplitudes of 5 to 40 nm, based on their liquid or gel nature. For the evaluations, measurement files were imported by custom evaluation software (Mathematica 13.2, Wolfram) and the procedure was performed as described in the Results section to achieve *G*’, and *G*”. For η, *G*” for all droplets of the same dataset were fitted together linearly, the uncertainty is the error of the fit. In isolated cases, due to experimental uncertainties such as phase noise for phase values close to zero and particle spring constants significantly larger than the cantilever spring constant, *G*’ and *G*’’ can obtain unphysical, negative values (i.e., phase < 0° or > 90°), which were discarded during evaluations. The radius of condensates was measured using ImageJ from the phase contrast images.

To resolve the liquid to gel transition and the subsequent rejuvenation of the condensates, we took advantage of the salt concentration sensitivity of the droplets. After the droplets were let to settle for 2 h and measured, the supernatant was exchanged to 0.65 M KCl concentration. Then, the same ten condensates were probed and the concentration in the buffer was once again reduced to 0.5 M KCl. Afterwards, the supernatant was carefully exchanged back to 0.65 and 1 M, and the condensates were measured. To temporally resolve the gelation without disturbing the condensates, we increased the humidity in the chamber by temperature. 300 µl water was added in the chamber and the dish was heated to 55°C and kept at this temperature for 70 min to allow the evaporated water to slowly condense on the bulk drop, reducing the KCl concentration of the dilute phase.

For crosslinking the condensates to change their properties from liquid to gel, droplets were let to settle for about 2 h until most coalescence events have ceased. Afterwards, formaldehyde (252549) was added to the bulk material to a concentration of 4%. The pK in the dilute phase of the bulk also crosslinks, which may cause contamination of the cantilever. Therefore, the supernatant was exchanged with the supernatant of another pK-H droplet after 2 h of incubation. This avoids cantilever contamination while preserving the composition of the dilute phase and verifies that a change in the supernatant does not cause the solidification of droplets if the concentration stays equivalent.

### 4.3. Flat amplitude deflection calibration

During SPM experiments, the cantilever is subject to hydrodynamic drag force artifacts due to viscous friction of the cantilever with the liquid, resulting in a phase lag [73]. To determine this, the indenter probe was landed on the surface of the polystyrene dish and the head height was harmonically driven at low amplitudes of 2 nm. A preload of 200 pN on a hemispheric Si_3_N_4_ (elastic modulus, *E* ∼ 200 GPa) indenter probe with a 5.13 µm radius of curvature contacting a flat polystyrene (*E* ∼ 3 GPa) surface leads to a total deformation of less than 0.01 nm for the probe and the dish surface according to Hertzian contact mechanics. This is negligible compared to the 2 nm deflection of the cantilever. Hence, it can be assumed that the head height oscillations and the cantilever deflection are identical.

Both the driven head height oscillation *h*(*t*) and the resulting deflection oscillation *F*(*t*) are fitted with Eq.1 and 2. The phase lag at each frequency is calculated as

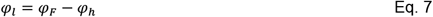

and the resulting data is well described by the linear model

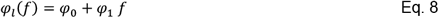

Where *φ*_0_ and *φ*_1_ are fit parameters.

### 4.4. Surface energy as a spring

To calculate the contribution of the surface energy to the droplet mechanics, its expression as a spring is necessary. In the undistorted condition, the volume of the particle is modeled as that of a sphere with radius *R*_2_, resulting in a volume of 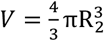. Upon indentation, the shape can be approximated by an oblate spheroid with identical volume (Poisson’s ratio 0.5) of 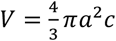, where *a* and *c* are the long and short semi-axis, respectively. Thus, the long semi-axis as a function of the short semi-axis at constant volume is given by 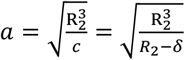 where δ is the deformation (i.e., indentation). The surface area *S* of an oblate spheroid is given by 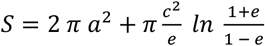 with 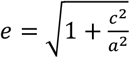. Thus, the surface energy of a spheroid is given by *E* = *γS* where *γ* is the surface energy density. With 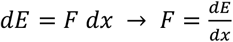 and with Hooke’s law, 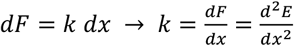, the spring model resulting from surface energy changes due to spheroidal distortion is given by

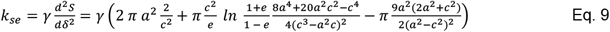

and thus

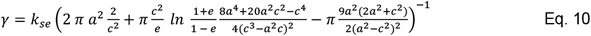

### 4.5. Hertzian contact model as a spring

During SPM measurements, the spring constant of the droplet is determined. The Hertzian contact model is necessary to convert between the complex spring constant and the complex shear modulus. Here, the hemispheric morphology of the condensates allows us to neglect the interaction between droplet and sample dish and by that the use of single Hertzian model. We also employed the double Hertzian model [70], which can be utilized for spheric droplets (Supplementary Information).

The contact model between an infinitely hard sphere (the indenter, subscript 1) and a soft sphere (the condensate, subscript 2) is given by

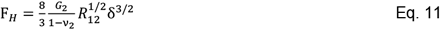

with

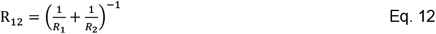

where *F*_H_ is the applied force, δ is the indentation, ν_2_ is Poisson’s ratio of the particle, *G*_2_ is the shear modulus of the particle, and *R*_1_ and *R*_2_ are the radii of curvature of the indenter and the particle, respectively. Using Hooke’s law, 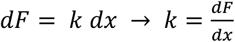, the spring given from the indentation of a viscoelastic material is

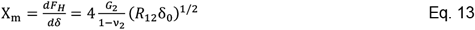

where δ_0_ is the preload indentation. Thus,

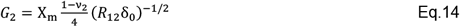

follows.

### 4.6. Fluorescent recovery after photobleaching (FRAP)

For FRAP experiments, 5% FITC labeled pK (P3543) was mixed with the unlabeled counterpart. The rest of the preparation was performed as stated above. Experiments were performed with a Nikon Ti2 Eclipse microscope and a 20x/0.75 objective. A circular area with a radius of 1.25 µm was bleached on droplets 2-3 times the bleach spot radius for 1 s at a laser power of 250µW and 488 nm wavelength followed by a 2 min acquisition of the recovery. As the internal arrangement of the molecules inside the droplet is faster than the exchange with the surrounding environment, the recovery was related to the movement of the molecules from the unbleached region within the droplet [13]. The FRAP data were normalized to the intensity of a reference unbleached droplet to account for photodecay from imaging and loss of fluorescence. Afterwards, the corrected data were fitted for deriving the diffusion coefficient using the half-time of recovery [74, 75]. Here, the normalized fluorescence intensity *F*_norm_ as a function of time *t*, where *t* = 0 s corresponds to the first measured point after the bleaching pulse, is calculated from the fluorescence intensity of the region of interest *F*_roi_, the fluorescence intensity of the reference area *F*_ref_, and the background fluorescence intensity *F*_bkgd_ as

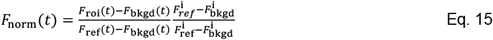

Where *F*^i^ denotes the time average of the fluorescence intensities for *t* < 0 s. *F*_norm_ is subsequently fitted with

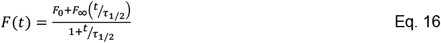

where *F*_0_ is the normalized fluorescence intensity at *t* = 0 s, *F*_∞_ is the fluorescence intensity at *t* → ∞, and τ _1/2_ is the half-time of recovery. From τ _1/2_, the diffusion coefficient *D* can be calculated as

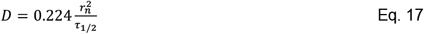

where *r*_n_ = 1.25 µm is the nominal radius of the bleach spot. Furthermore, the viscosity η can be calculated from *D* according to [8]

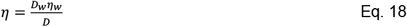

based on the diffusion coefficient *D*_w_ = 110×10^−12^ m^2^ s^-1^ and the viscosity η_w_ = 890 µPa s of the dye in water derived for globular proteins [8].

## Data and Code Availability Statement

The data that support the findings of this study and the custom codes in Wolfram Mathematica for analyzing primary data on rheological properties are available on Github. https://github.com/TheMashaghiLab/Publication_Biomolecular_Condensate_Rheology

## Author Contributions

Conceptualization: A.N., A.M. Methodology: A.N., O.A., A.M. Investigation: A.N. Software: O.A. Visualization: A.N. Formal Analysis: A.N., O.A. Writing & Draft Preparation: A.N. Writing-Review & Editing: A.N., O.A., A.M, Project Administration, Funding acquisition & Supervision: A.M.

## Acknowledgments

The authors would like to thank Kostas Tassis for assistance with confocal microscopy and FRAP measurements, as well as Liru Feng from Biological & Soft Matter, Leiden University for fruitful discussions. We are also grateful to the technical support of Vahid Sheikhhassani from Medical Systems Biophysics and Bioengineering Laboratory, Leiden University as well as Sebastian Friedrich and Andre Koernig from Bruker.

