## Supplementary Information for "Scanning probe microscopy elucidates gelation and rejuvenation of biomolecular condensates"

This file includes:

Supplementary Figures S1 to 7

Supplementary Text with the description of indentation determination for droplets after gelation and double Hertzian contact model

Other Supplementary Information for this manuscript include Supplementary Movies 1 and 2.

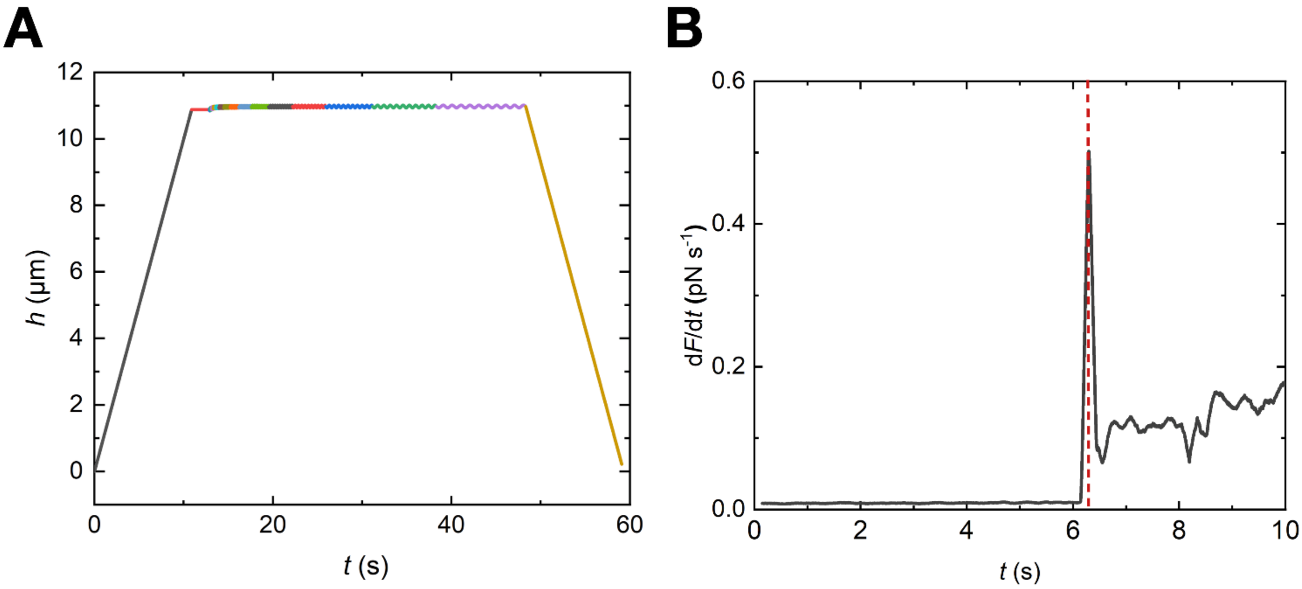


**Supplementary Figure S 1: A)** A representative curve of head height over time *h*(*t*), corresponding to the cantilever deflection force over time *F*(*t*) depicted in Figure 1B. **B)** Exemplary numerical derivative of the approach segment of *F*(*t*) signal depicting a sudden change of force to appear as a peak in the derivate data allowing the determination of the approximate contact time between the cantilever and the condensate.

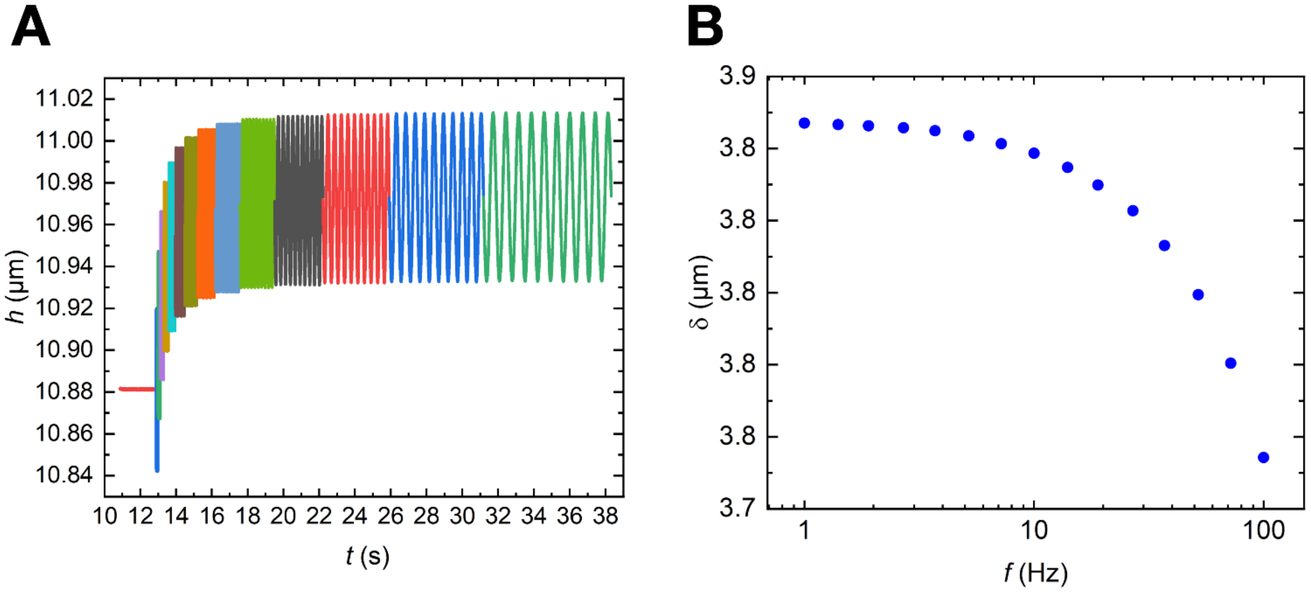


**Supplementary Figure S 2***:* **A)** A representative curve of the modulation segment of head height over time *h*(*t*) depicting the change in the indentation with time which may originate form e.g. the temperature drift of the cantilever or the creep of the droplets. **B)** Indentation depth δ calculated separately as the time average of *h*(*t*) for each frequency to account for the change in indentation over time.

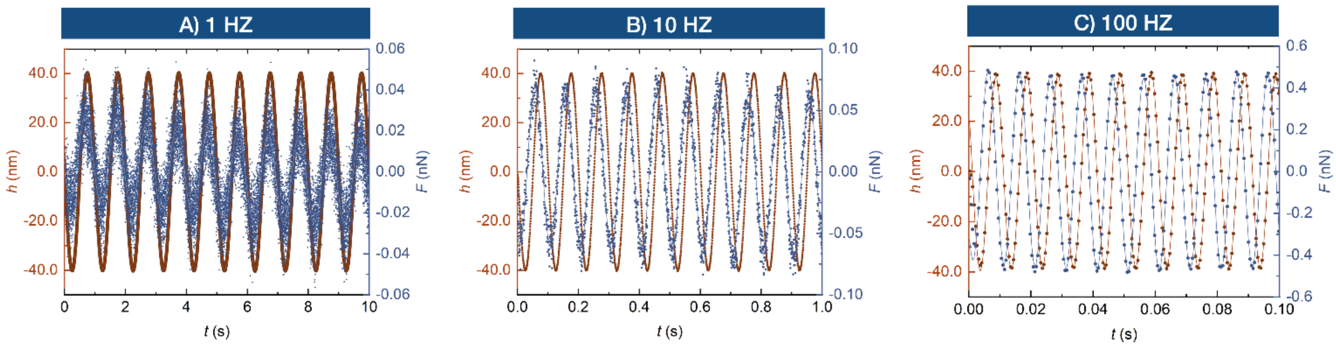


**Supplementary Figure S 3:** Experimental data points and the fit of *F*(*t*) and *h*(*t*) at **A)** 1 Hz **B)** 10 Hz **C)** 100 Hz demonstrating the strong frequency-dependent response of the liquid droplets.

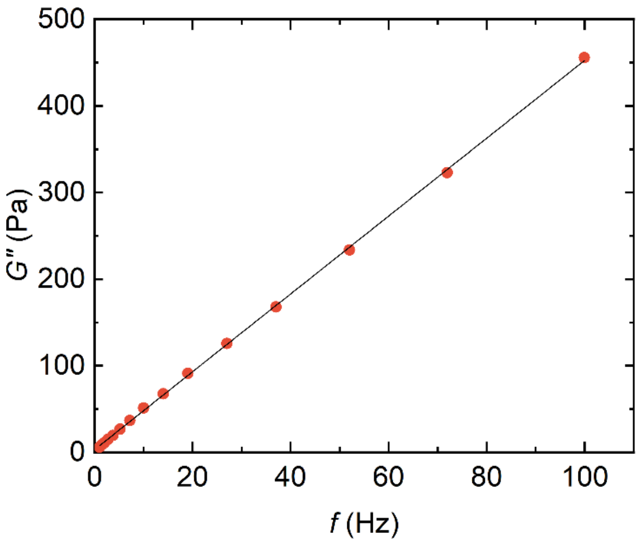


**Supplementary Figure S 4:** An exemplary linear fit of the of *G*" for determination of viscosity leading to (4.49 ± 0.02) Pa s.

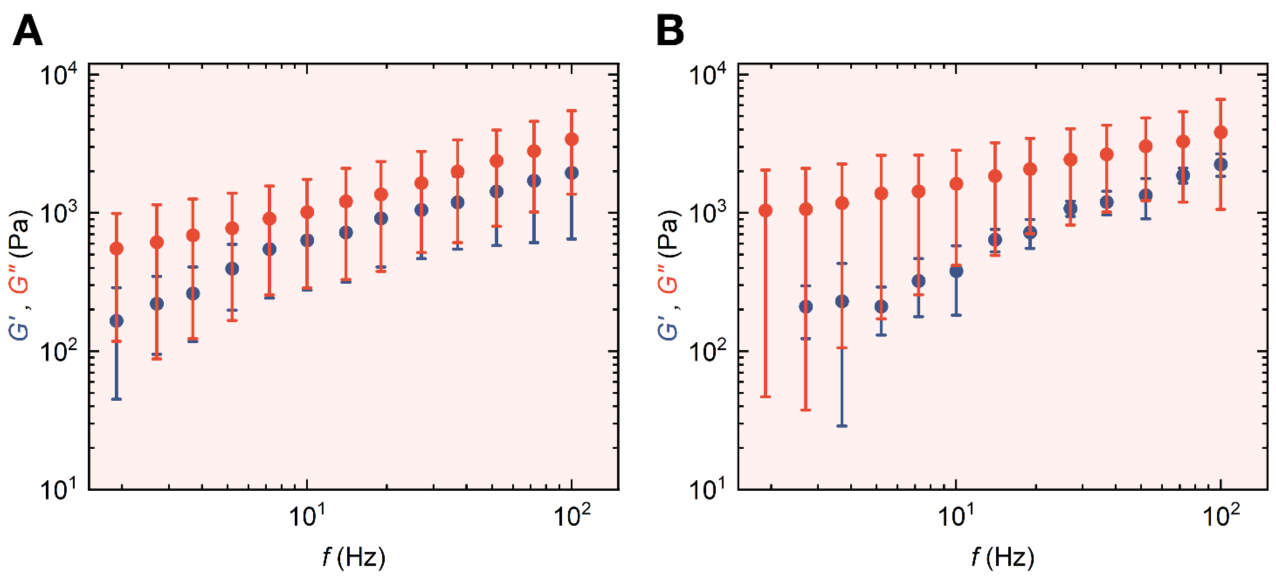


**Supplementary Figure S 5:** Elastic (*G*', blue) and viscous (*G*", red) moduli of pK-H condensates formed at 1 M KCl and **A)** Measured after dilution of supernatant to 0.65 M KCl. **D)** Measured after exchange of supernatant back to 0.65 M KCl, after gelation at 0.5 M (mean ± SD, $N = 10$).

**Indentation determination of droplets after gelation**

Supplementary Figure S 6 shows the deflection signal of a typical approach for droplets in their liquid form and after transition to gel-like. The description for the indentation determination of liquid condensates can be found in the main text chapter 2.1. For the droplets after gelation, an approach generally used in SMP measurements is employed. With the probe at a distance from the sample surface, the force is constant and close to zero. As soon as the probe contacts the sample, the force shows a roughly linear increase in force. In order to determine the time of zero indentation $t_0$, a region in the flat section of ca. 1 s duration just before the point of contact and the longest linear portion directly after $t_0$ are specified. Both regions are linearly fitted and $t_0$ is calculated as the intersection of the two fitted lines. From $t_0$, the head height of zero indentation can be read from $h_r(t_0) = h_0$.

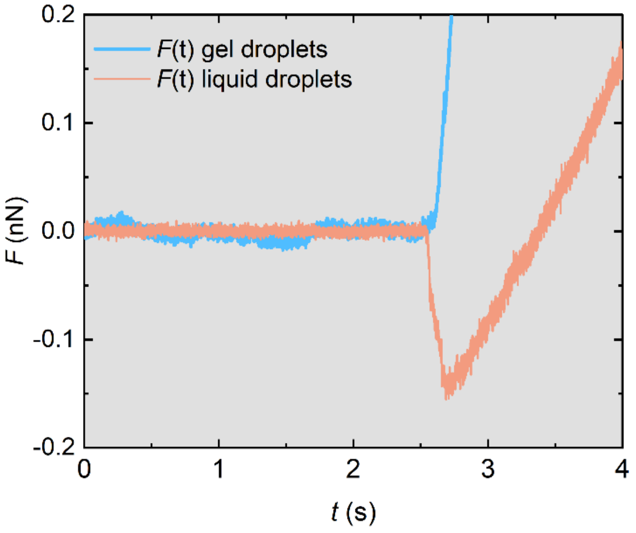


**Supplementary Figure S 6:** Representative curve of the approach segment of cantilever deflection force over time, $F(t)$ for liquid droplets in pink and droplets after gelation in blue.

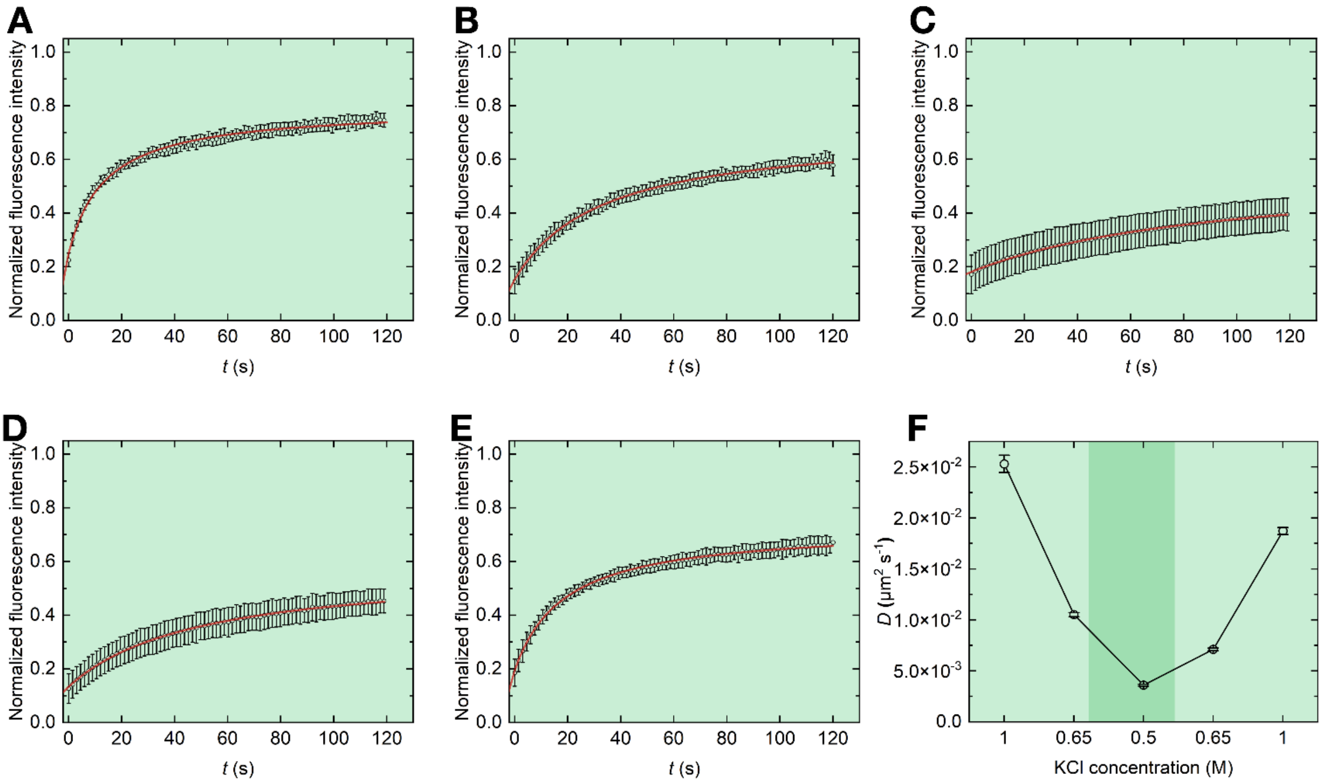


**Supplementary Figure S 7:** Time-dependent normalized fluorescence intensity depicting the recovery of the bleached area (circular data points) as well as the fit with Eq. 16, for condensates formed at 1 M KCl and **A)** Measured at 1 M KCl **B)** Measured after dilution of supernatant to 0.65 M KCl. **C)** Measured after dilution of supernatant to 0.5 M KCl. **D)** Measured after exchange of supernatant back to 0.65 M KCl. **E)** Measured after exchange of supernatant back to 1 M KCl **F)** The corresponding diffusion constant ($D$).

**Double Hertzian contact model as a spring**

For spherical particles, where the indenter-particle and the particle-substrate contact areas are comparable it is crucial to consider the deformation on both contact sides of the condensates. The double Hertzian contact between an infinitely hard sphere (the indenter, subscript 1), a soft sphere (the condensate, subscript 2) and an infinitely hard flat surface was derived as [70]

$$\mathrm{F}_{DH} = \frac{8}{3}\frac{G_2}{1-\nu_2}R_{12}^{1/2}(r\,\delta)^{3/2}$$

with

$$\mathrm{r} = \frac{R_2^{1/3}}{R_{12}^{1/3} + R_2^{1/3}}$$

$$\mathrm{R}_{12} = \left(\frac{1}{R_1} + \frac{1}{R_2}\right)^{-1}$$

where $F_{DH}$ is the applied force, $\delta$ is the indentation, $\nu_2$ is Poisson's ratio of the particle, $G_2$ is the shear modulus of the particle, and $R_1$ and $R_2$ are the radii of curvature of the indenter and the particle, respectively. Using Hooke's law, $dF = k\,dx \;\rightarrow\; k = \frac{dF}{dx}$, the spring given from the deformation of a viscoelastic material

$$\mathrm{X_m} = \frac{dF_{DH}}{d\delta} = 4\frac{G_2}{1-\nu_2}(R_{12}\delta_0)^{1/2}r^{3/2}$$

where $\delta_0$ is the preload indentation. Thus,

$$G_2 = \mathrm{X_m}\frac{1-\nu_2}{4}(R_{12}\delta_0)^{-1/2}r^{-3/2}$$

follows.
